# Robust dynamical invariants in sequential neural activity

**DOI:** 10.1101/379909

**Authors:** Irene Elices, Rafael Levi, David Arroyo, Francisco B. Rodriguez, Pablo Varona

## Abstract

By studying different sources of temporal variability in central pattern generator circuits, in this paper we unveil distinct aspects of the instantaneous balance between flexibility and robustness in sequential dynamics –a property that characterizes many systems that display neural rhythms. The level of irregularity and coordination was characterized using intrinsic time references and intervals in long recordings of the pyloric central pattern generator. The analysis demonstrated strong robustness of transient dynamics in keeping not only the activation sequences but also specific cycle-by-cycle temporal relationships in the form of dynamical invariants. The rich dynamics of neurons and connections balance flexibility and coordination to readily negotiate the interactions between neurons and produce the resultant rhythm. In particular, two dynamical invariants were identified between time intervals that build the sequence, existing even outside steady states. We suggest that invariant temporal sequence relationships could be present in other networks, including those related to brain rhythms, and underlie rhythm programming and functionality.

## 1 Introduction

Robust sequences of neural activations can be found in any nervous system, from simple invertebrate circuits^1–4^ to almost any vertebrate system.^5–12^ As both experimental and theoretical studies show, temporal sequence generation is a key computational phenomenon to encode, control and execute information in sensory, central and motor networks.^13–18^ In many cases, robust sequences underlie what is usually simply viewed and termed as a rhythm, a spatio-temporal pattern, or alternating patterned activity. Unveiling general principles in the generation and coordination of neural sequences, particularly in transient regimes, is an important step in relating neural activity to function and has potential impact in other fields such as rehabilitation technology, robotics and control theory.^19–22^

Central pattern generators (CPGs) are neural circuits that produce flexible rhythmic motor patterns.^23,24^ The robust and highly coordinated neuron activation sequences underlying their rhythms arise from the combination of intrinsic membrane currents and properties of the synaptic connections.^25,26^ Invertebrate CPGs are key neural circuits to understand rhythm generation and coordination, as their cells and connections have been identified and mapped, like in the crustacean pyloric CPG that we use in this study.^25,27–29^ Most CPGs have what is called a non-open topology,^30^ i.e., all neurons in the CPG receive input from other cells in the circuit for transient closed-loop computation. An essential component of these sequence generator networks is reciprocal inhibition between pairs of neurons. Mutual inhibition together with electrical coupling and other non-reciprocal interactions^30,31^ underly the timing of neuron activations that shape each cycle.^24,32,34^ CPGs activate the muscles that produce motor rhythms (in the case of the pylorus, a triphasic rhythm) and their flawless function is crucial for the animal survival.^27,28^

Experimental and computational studies of CPGs traditionally examine their rhythmic output from periodic spiking-bursting regimes. Recordings of crustacean pyloric CPG neurons in the semi-intact network typically show regular sequential behavior of the circuit, e.g.^25,35^ On the other hand, *in vivo* recordings of CPG activity display further temporal characteristics and a larger degree of irregularity than *in vitro* recordings.^35–37^ Recordings of isolated cells show that most of them are highly irregular.^38–41^ Rich intrinsic cell and synaptic dynamics, arising from different time scales,^26^enable neurons comprising the CPG to readily establish a network rhythm in concert.^25^Recent work has proposed that variability in cycle period can be controlled by synaptic feedback.^42,43^ External and intrinsic factors continuously effect the system inducing transients and therefore making them an important element of the system functionality, which can only be observed in recordings of irregular activity.

Early on, the CPG community identified the importance of phase maintenance for motor function.^44–46^ Other studies in the pyloric CPG have reported approximate maintenance of phase-frequency relationship when altering the rhythm speed by current injection^33,47,49^ or by temperature changes.^35,50^ By quantifying average delays and periods and comparing across preparations, authors could show that certain elements of the rhythm maintain relative timing with changes in frequency.^35,47,48^ In this context, most works discard irregular activity, as well as, transient changes. Information regarding variability cannot be ignored when characterizing the instantaneous generation and coordination of neural sequences. This information is lost under traditional average analyses.

Here, we address the changes in a cycle by cycle ongoing CPG rhythm, and argue that all neurons participate in the production and tunning of the system, so that each contributes to the instantaneous negotiation of the resultant pattern. We show that the flexibility from rich intrinsic neuron dynamics is actively bounded by the connectivity in a cycle-by-cycle negotiation that produces coordinated sequential activity, even during transients. Thus, to expose this process we also studied irregular rhythms, i.e., activity that presented high variability in period, phase between neurons, or burst duration in the same experiment. In particular, we used two sources of rhythm irregularity: intrinsic variability in the preparation, and irregularity induced by ethanol. Our departing hypothesis is that the analysis of non-periodic regimes can unveil properties of the underlying robust dynamics controlling rhythm coordination, which remain unnoticed in regular rhythm regimes. In spite of the large variability seen in the experiments, we report two robust *dynamical invariants* in the form of linear correlations between time intervals that build the sequence. These invariants were strongly preserved under different conditions and were found using a time reference frame that took into account the asymmetric topology of the system. We characterized these invariants and related them to the underlying balance of influences between the rich intrinsic dynamics of neurons and the asymmetric connectivity of the circuit. We hypothesize that dynamical invariants participate in the instantaneous coordination of the different muscles innervated by the CPG neurons, and therefore can be linked to the efficient cycle-by-cycle performance of motor activity of the system in different circumstances.

Considering regular bursting activity and a dynamic-clamp protocol that altered a CPG synapse, previous work reported a dynamical invariant in which the ratio between the resulting change in average burst duration and the change in average phase lag between PD and LP neurons was tightly preserved in all preparations.^51^Here we follow this terminology, but we refer to robustly preserved instantaneous interval relationships within the variability of a preparation, as opposed to cross-preparation averaged phase maintenance. We believe this term better reflects the transient information exchange in the circuit.

Beyond CPGs, dynamical invariants might be present in a wide variety of circuits throughout the nervous system. Frequency independent temporal ordering has been observed in different neural systems, including the hippocampus and the cortex.^52,53^ The study of instantaneously preserved temporal relationships in brain rhythms can provide key insights regarding their functional role in the context of precise sequential information encoding and execution. We argue that the insight gained from examining irregular activity transients and dynamical invariants in simple CPG circuits will lead to deeper understanding of robust sequential activations in functional brain rhythms.

## 2 Results

### 2.1 Characterization of variability of spiking-bursting activity in CPGs

The pyloric CPG of the crustacean stomatogastric nervous system presents a characteristic rhythm with three main components in a robust sequence: the Lateral Pyloric (LP) neuron, a group of six to eight pyloric neurons (PY), two electrically coupled Pyloric Dilator (PD) neurons and the Anterior Burster (AB), also electrically coupled to the PDs.^23,27,54^ Panel A in ***Figure 1*** shows an example of extracellular recording of the LVn nerve in which these three components can be clearly distinguished, along with intracellular recordings from PD and LP neurons. In control conditions, this circuit typically produces a regular and robust rhythm with nearly constant burst durations and hyperpolarization intervals *(**Figure 1***, panel A).

**Figure 1:**
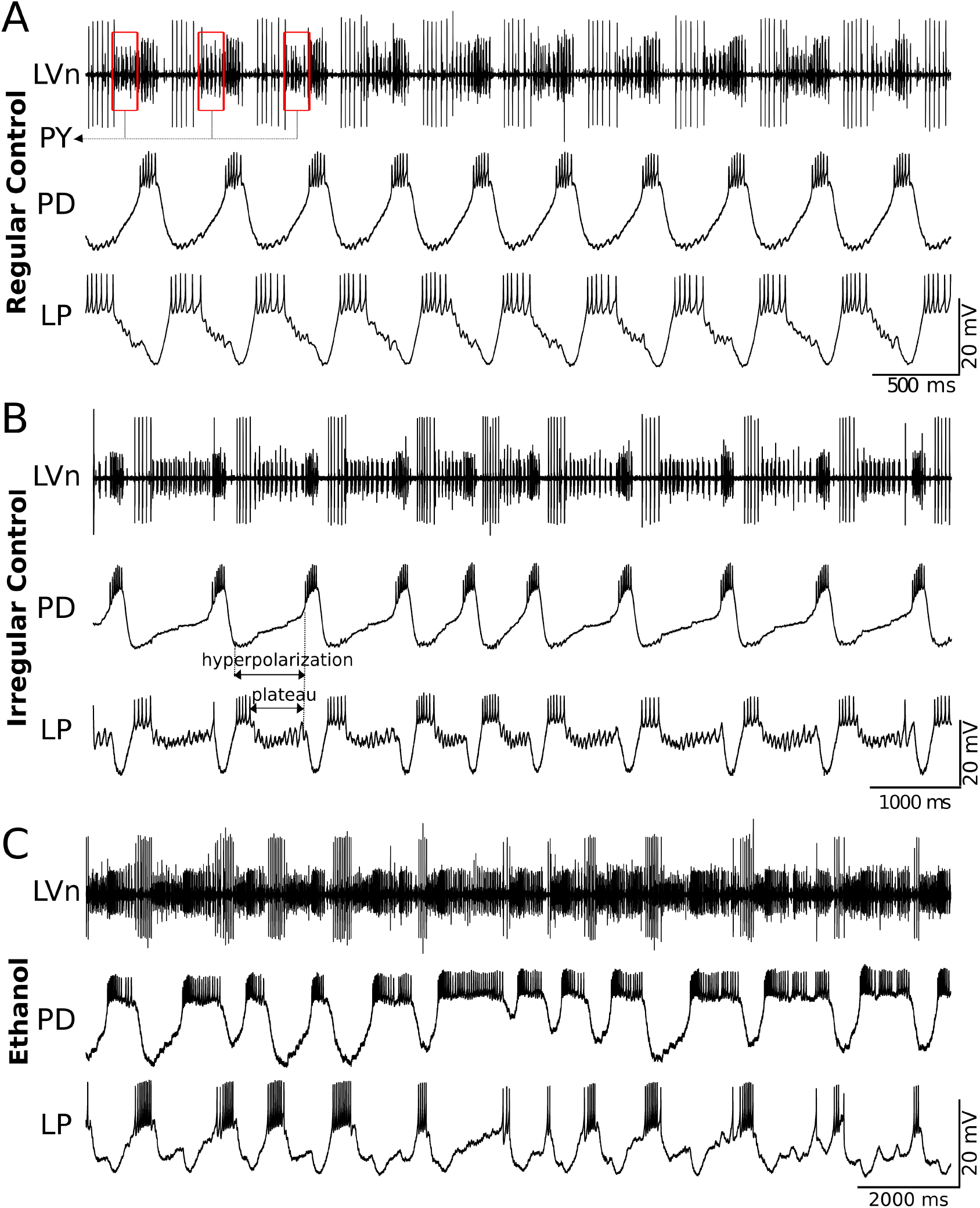
Examples of sequential activity produced by the pyloric CPG. The traces correspond to simultaneous extracellular recordings of the LVn nerve (upper trace) and intracellular recordings of PD and LP neurons in the intact CPG. Panel A: An example of the characteristic regular triphasic spiking-bursting activity in this CPG circuit. Large spikes in the LVn trace correspond to the LP neuron. Note that LP spikes occur in antiphase with PD spikes and the respective IPSPs can be observed in the PD neuron trace. PY spikes can be observed in the extracellular recording after the LP and before the PD spikes (red boxes in the upper trace). PD and LP burst durations and hyperpolarization intervals are nearly constant in the recordings. Panel B: Example of transient irregular spiking-bursting activity in control conditions. Note the irregular hyperpolarizations and variability in LP plateaus as compared to the regular trace shown in regular control conditions. Panel C: Example of irregular spiking-bursting activity with addition of ethanol (170 *mM*). Note that PD neuron presents flexible and long burst durations while LP neuron has more restricted burst durations. In all cases the sequence of neuron activations LP-PY-PD is preserved, in spite of the intrinsic or induced irregularity.

Irregular activity can also be seen in cases of intrinsic variability in the preparation, which may be a result of the external neural modulation, slightly damaged neurons or severed modulator nerves, etc. Panel B in ***Figure 1*** shows an example of extracellular (LVn) and intracellular recordings from PD and LP neurons in a stomatogastric ganglion with intrinsic variability. There are clear differences from the regular activity recordings shown in panel A: hyperpolarizations of both neurons are irregular and, while PD bursts remain more constant, LP presents longer plateaus and higher variability in burst duration. Irregularity can also be induced chemically, e.g. evoked by application of ethanol (***Figure 1***, panel C). Pyloric rhythm under ethanol is characterized by a remarkably flexible and long PD burst duration, and variability in hyperpolarization in both neurons. At the same time, LP burst duration presents much less variability as compared to the PD neuron. Despite the large irregularity induced by ethanol, the sequence LP-PY-PD in the rhythm is still preserved. After washing or ethanol evaporation, regular activity is recovered.

The analysis of variability of CPG rhythms in all conditions was assessed by defining specific intervals using precise time references as: the first and last spike of a burst from intracellular recordings. We chose six intervals (defined in ***Figure 2***): *Period, LPPD delay*(corresponding to PY neuron activity), *LPPD interval*, PD burst duration *BD_PD_*, LP burst duration *BD_LP_* and *PDLP delay*, and studied variability in long recordings using their coefficient of variation (*C_v_* = *σ/μ* · 100(%)).

**Figure 2:**
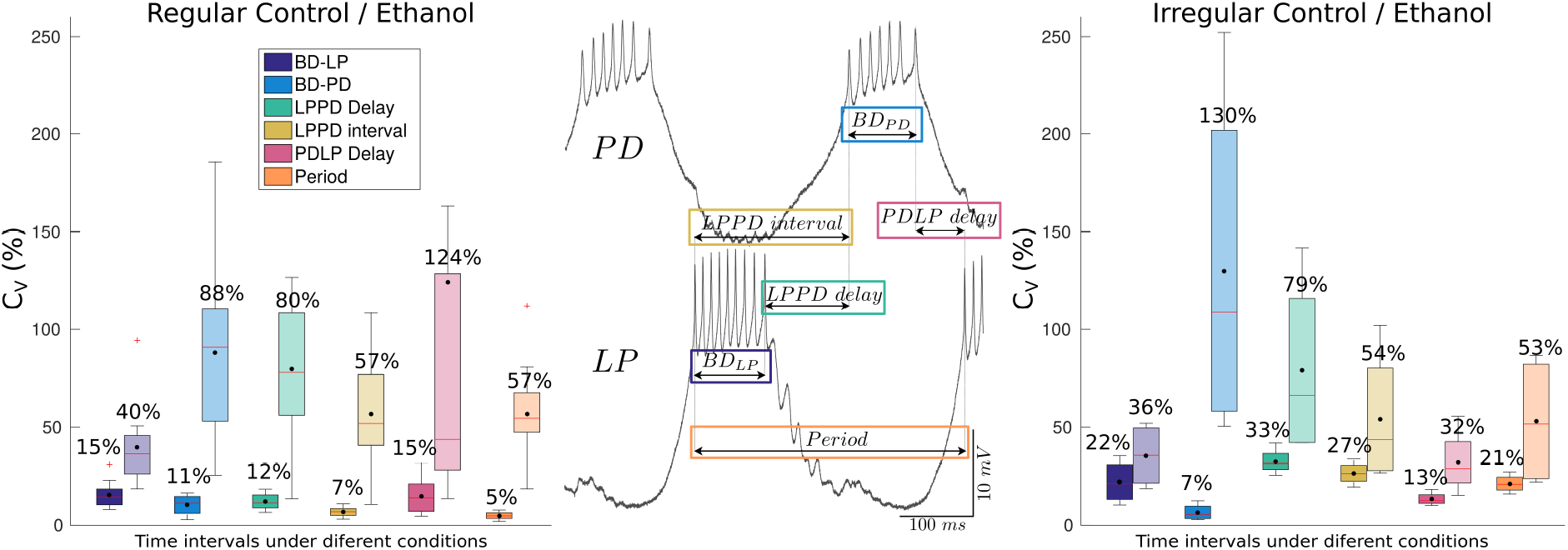
Definition and variability analysis of temporal intervals considered in this study to characterize the CPG cycle-by-cycle rhythm. Central panel: Scheme of the definition of the measured time intervals. *BD_PD_* and *BD_LP_*: burst duration defined as the time interval from the first spike to the last spike of PD and LP neurons, respectively; *LPPD delay:* interval defined from the LP last spike to the PD first spike; *LPPD interval:* interval defined from the LP first spike to the PD first spike; *PDLP delay:* interval defined from the PD last spike to the next first LP of the following burst; *Period:* interval from the LP first spike to the next LP first spike in the following burst. Left and right panels: Boxplots of the coefficient of variation for the six measures in control conditions (darker color) and under the influence of ethanol (lighter hue boxes). Mean values (black dots) are displayed on top of each box. Left panel: Quantification of the variability in long recordings for preparations that were regular in control conditions (n=12). The coefficients of variation are small (5— 15%) in control conditions. Under the influence of ethanol, in lighter colored boxes, there is a large increase in variability for *BD_PD_* (88%), *LPPD delay* (80%) and *PDLP delay* (124%) while *BD_LP_* is more restricted in variability (40%). Right panel: Intrinsically irregular preparations (n=4). One can observe an increase in variability of *LPPD delay* and *LPPD interval* due to the irregular hyperpolarization intervals in control conditions (see *Figure* 1). After applying ethanol, there is even larger variability in *BD_PD_* (130%) and *LPPD delay* (79%) while *BD_LP_* variability remains more restricted (36%).

Note that some of these intervals are different from those used in other pyloric CPG studies that consider as time reference the beginning of the PD burst. In most studies, the PD neuron burst beginning is used as the time reference for cycle period and delays of the other so-called follower neurons.^47,55,56^ In addition, a considerable variability across individual preparations has been observed in phase-frequency relationships when the pacemaker group is used as the time reference.^56,57^ Since PD neurons have strong inertia from electrical coupling among all cells in the pacemaker group, other time reference frames are more suitable to address the balance between flexibility and robustness, see also.^48^

***Figure 2*** compares the average of the coefficient of variation of the considered intervals, described in the central panel, for preparations with departing regular (left panel) and irregular (right panel) activity in control conditions and under ethanol by means of box-plots. Boxes in darker color correspond to control conditions. In regular control preparations, the values the *C_v_* of the six intervals went from 5% to 15%, the highest corresponding to *PDLP delay* and *BD_LP_*. In the case of intrinsic irregular activity, the variability increased in all intervals except for *BD_PD_*. One can observe that in control conditions the variability was small but still leaving room for flexibility. Under the influence of ethanol (lighter hue boxes), both regular and irregular preparations increased the variability in all intervals. In particular, PD burst duration *BD_PD_* (88%— 130%) and the *LPPD delay* (80%—79%) presented a larger variability while *BD_LP_* was lower (40%— 36%). Note that these three intervals build the triphasic rhythm. The interquartile range of the boxes indicates the variability among preparations and in the case of ethanol highlights the differences of its effect on the rhythm. Overall, the system showed a wide range of variability specific to the distinct intervals that shape the rhythm with large variability in some, such as *BD_PD_*, and smaller variability in others (e.g., *BD_LP_*).

### 2.2 Dynamical invariants

In order to identify factors shaping the CPG transient rhythm negotiation, i.e., the process of balancing flexibility and robustness of timings and sequence, we analyzed the cycle-by-cycle intervals defined above in regular and irregular rhythms. The underlying question is whether there is any property or temporal relationship in the rhythm, in addition to the sequence of neuron activations, which is preserved under different conditions (regular rhythms, intrinsic irregularity or ethanol), i.e., a *dynamical invariant*.

Departing from well-defined time references at the burst beginning and end in the LP and PD neurons, we analyzed *Period, LPPD delay*, *LPPD interval, PDLP delay* and burst durations *BD_PD,LP_*, and searched for preserved correlations between pairs of intervals, even when the rhythm was very irregular. We performed this analysis cycle by cycle in long continuous intracellular recordings. It is important to note, that most relationships between intervals were not preserved, such as *BD_PD,LP_* as a function of the *Period, BD_LP_* or *LPPD interval* as a function of *BD_PD_* or *BD_PD,LP_* as a function of the *LPPD delay* (see ***Table* 1**), and none of the intervals defined with the PD time reference frame. However, we found two relationships that presented strong linear correlations in all preparations: the measured *LPPD delay* and *Period* and *LPPD interval* and *Period* (see middle panel in ***Figure 2*** and ***Table* 1**). These dynamical invariants consistently remained tightly preserved with a slope close to one for many different preparations and under different conditions.

**Table 1:**
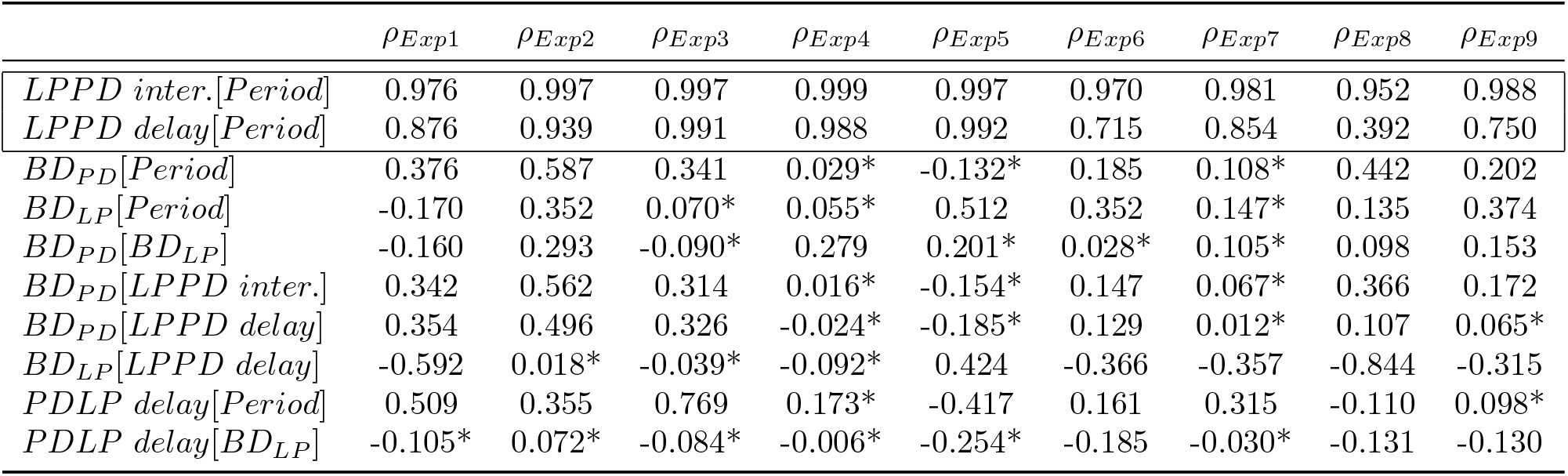
Values of the Pearson correlation coefficient *ρ* obtained for the different combinations of instantaneous intervals considered in this study for 9 representative experiments in control conditions (same preparations as in ***Figure 3*** and ***Figure 4***). Other experiments show similar results. Regression analysis indicated that both *LPPD interval* and *delay*(framed in the table) have a strong correlation with *Period* (*p* < 10^−7^ each) consistently in all the experiments, while other measured variables and not correlated. Note that the only intervals correlated to the Period are those defined in the LP time reference frame. * Slope not significantly different from 0 (*p* > 10^−7^).

***Figure 3*** depicts these two preserved relationships for 9 representative experiments in control conditions with their corresponding linear regression. The analysis includes both regular and intrinsically irregular rhythms (indicated with f). The linear regression shows that the ratio between the change from one cycle to the next in *LPPD interval, delay* and the change in *Period* is constant. The strong linear correlations, indicated the presence of these invariants despite the rhythm variability (*R*^2^ > 0.9 for *LPPD interval*[*Period*]). We also included the special case of experiment Exp. 8 in ***Figure 3*** with *ρ_Delay_*[*Period*] = 0.392. This low coefficient of correlation can be attributed to the very small variability in control conditions resulting from a remarkably fast and highly regular rhythm in this experiment, which hides the invariant relationship. Video 1 in the supplementary material illustrates, with the help of the sonification of the sequential activity, the presence of the invariants as the spiking-bursting activity progresses in time.

**Figure 3:**
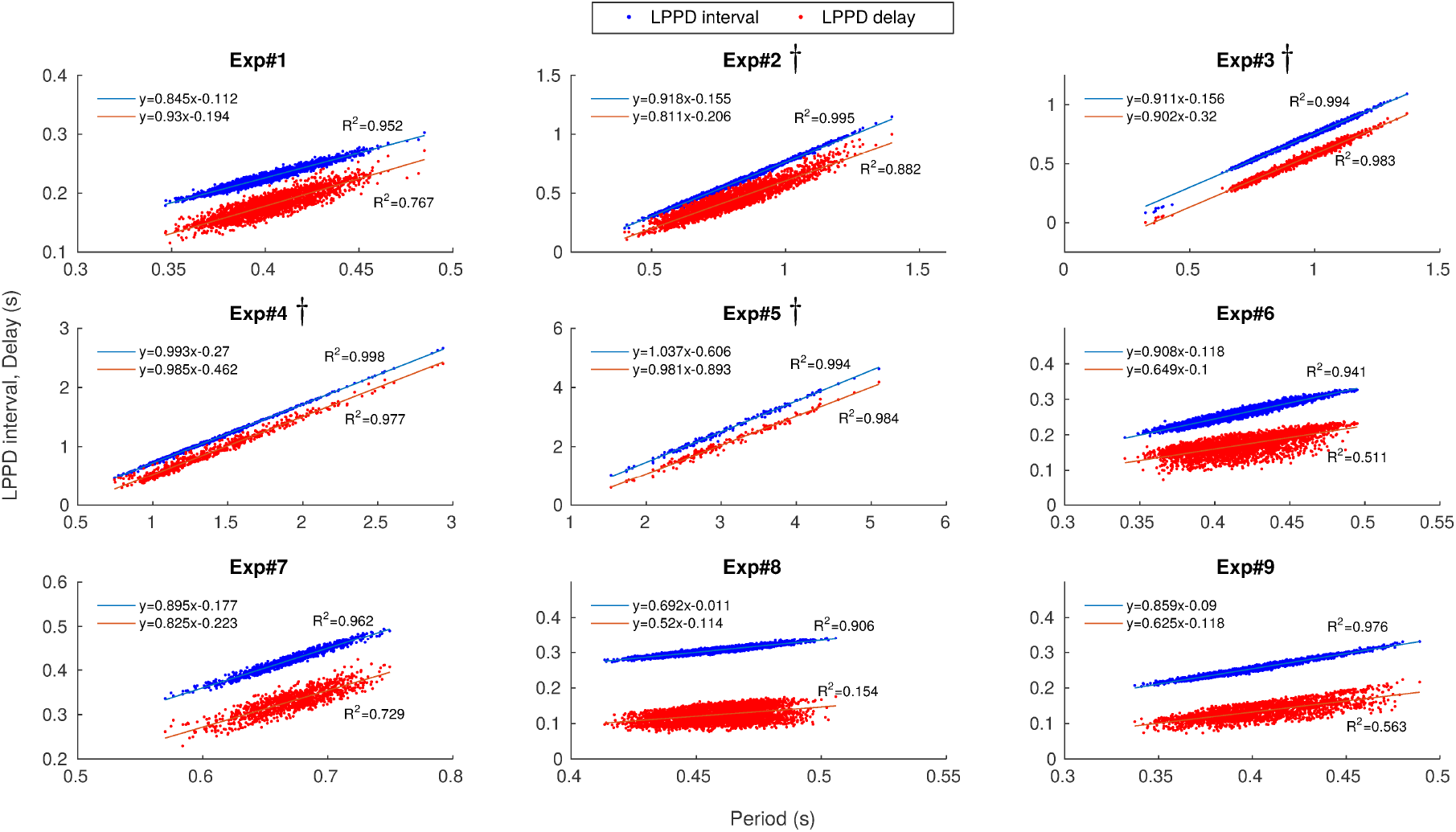
Presence of the two dynamical invariants in *control conditions* in 9 representative preparations. The correlation between *LPPD interval* and *Period* is shown in blue while the correlation between *LPPD delay* and *Period* is shown in red. Each point corresponds to one pyloric cycle of continuous recordings. Linear regressions are depicted for each experiment. Regression analysis showed that both LPPD interval and delay values increased with period (*p* < 10^−7^ each). The linear dependence is indicated by *R*^2^ values displayed for each experiment in the corresponding panel. *f* Intrinsically irregular preparations.

***Figure 4*** depicts these relationships for the same preparations illustrated in ***Figure 3*** under the influence of ethanol applied after control. Even in this condition, in which the variability of the measured intervals was very large, the invariants were still present. Note that now Exp. 8 yields *R*^2^ > 0.9 for both invariants. Under the influence of ethanol, the CPG rhythm can display very long bursts (lasting in some cases over 6 seconds). During some sections of the recordings in ethanol conditions, bursts in the sequence were lost. These sections that did not contained the required time references were removed from the statistics as illustrated in ***Suplementary figure* 3**. ***Suplementary table* 1** shows the percentage of dismissed burst which never exceeded 26% of the whole recording. The presence of the invariant in ethanol suggests that variability in *BD_PD,LP_* and variability in *LPPD delay* compensate each other cycle by cycle to sustain the invariants in the rhythm.

**Figure 4:**
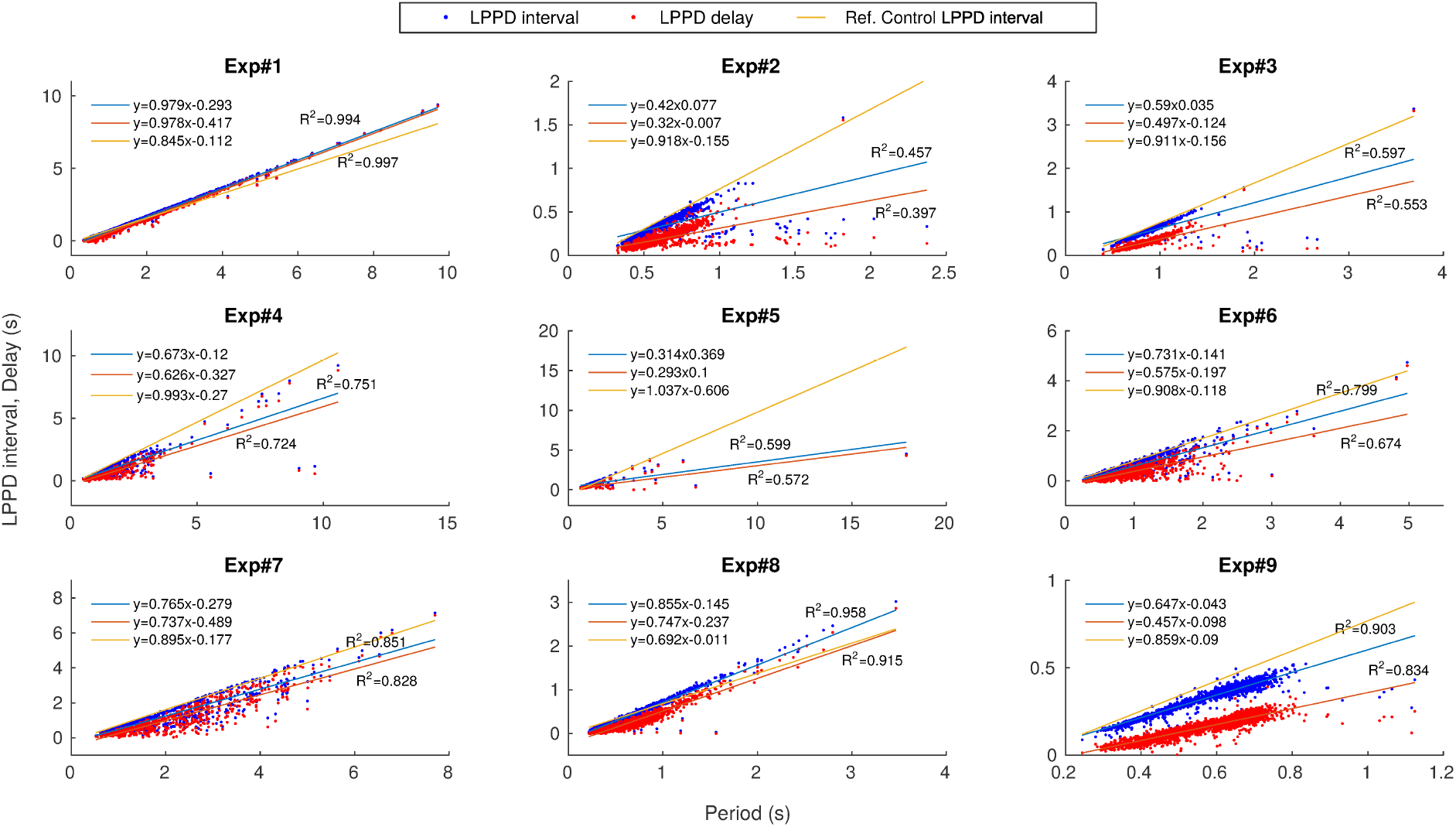
Presence of the two dynamical invariants under the influence of *ethanol* for the corresponding 9 preparations displayed in ***Figure 3***. The correlation between the measured *LPPD interval* and *Period* is shown in blue while the correlation between *LPPD delay* and *Period* is shown in red. Each point corresponds to one pyloric cycle. Linear regressions are depicted for each experiment. Regression analysis showed that both *LPPDinterval* and *delay* values increased with period (*p* < 10^−7^ each). The linear dependence is indicated by *R*^2^ values displayed for each experiment in the corresponding panel. Line in orange corresponds to the linear regression between *LPPD interval* and *Period* in control conditions shown in ***Figure 3***, and is provided to facilitate the comparison.

One notable property of the pyloric CPG network is its asymmetric inhibitory connectivity. This connectivity could play a key role in explaining the compensation process that creates the invariants, so that, if key synapses are removed, invariants should change or even disappear. Thus, we applied picrotoxin (PTX) 5 · 10^−7^ *M*, a glutamatergic synapse blocker, which blocked the fast inhibitory synapses (see ***Figure 5*** panel A). Panel B in ***Figure 5*** shows an example of PD and LP activity after applying PTX. One can observe the irregular shape of the LP bursting activity, allowed by the low PTX concentration, the absence of LP IPSPs in the PD trace and the removal of the LP plateau. A comparison of the coefficient of variation in three conditions, control, PTX and PTX+EtOH, is shown in panel C. In control conditions, the variability was small for all measures (5% – 15%). After applying PTX there was a slight increase in *C_v_* for all measures except for *LPPD delay* that reached 163%. Adding ethanol increased the variability even further (43% – 201%) with values similar to those obtained in experiments after applying ethanol alone (c.f.***Figure 2***) except for *BD_LP_* and *LPPD delay*, which showed larger variability after removing the connections from the PYs and AB with PTX.

**Figure 5:**
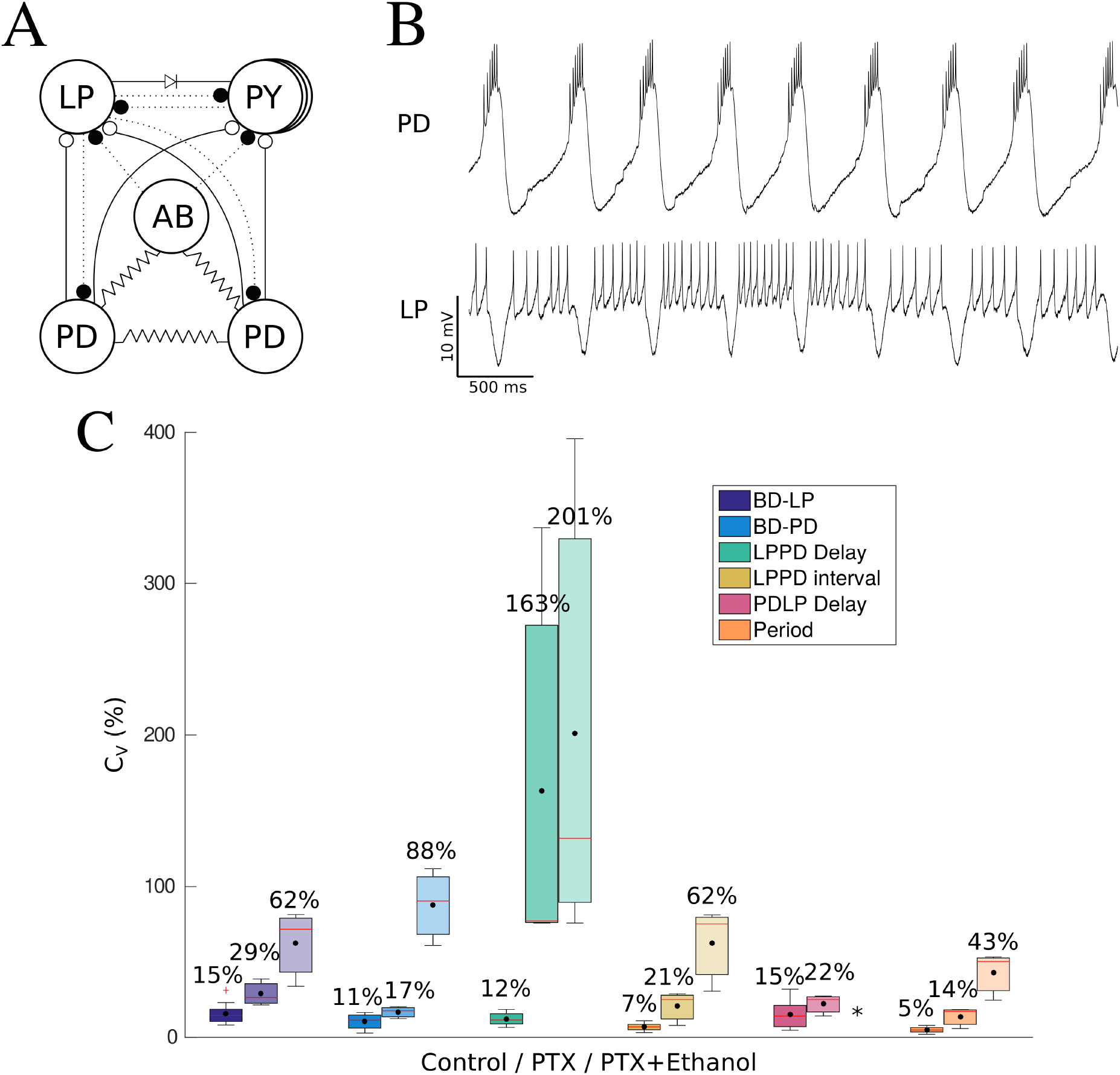
Results of blocking fast inhibitory synapses with *PTX*. Panel A: Scheme of the connectivity of the pyloric CPG after applying picrotoxin (PTX) 5 · 10^−7^ *M*. Dotted lines correspond to blocked fast inhibitory synapses. Panel B: Example of the spiking-bursting activity of the circuit after applying PTX. The traces correspond to simultaneous intracellular recordings of PD (upper trace) and LP (lower trace) neurons. Note that the characteristic IPSPs typical seen in the PD neuron trace are no longer present. Panel C: Coefficient of variation (*C_v_*) for the six measures in three conditions: control, first column for each measure (darkest color); after applying PTX 5 · 10^−7^ *M*, middle column; after adding ethanol to the PTX dilution, third column (lightest hue boxes). The highest variability in control conditions corresponds to *BD_LP_* and *PDLP delay* (15%), while after applying PTX the highest *C_v_* corresponds to *LPPD delay* with 163%, which is almost 14 times higher than in control. Variability in the other 5 measures also increases with PTX although more slightly. Adding ethanol to the PTX solution increases variability even further (43% – 201%).

***Figure 6*** shows the correlations in three experiments under three conditions: control, PTX, and PTX and ethanol. One can observe that the dynamical invariant *LPPD delay [Period]* was not preserved in the absence of fast synapses with slopes tending to 0 in all experiments, as opposed to the dynamical invariant *LPPD interval*[*Period*] that maintains a tendency similar to the corresponding control. A possible explanation for the preservation of this invariant is the LP burst duration *BD_LP_* variability, which manages to compensate PD variability, even under the effect of ethanol. More examples of the effect of PTX on the invariant correlations are provided in supplementary Fig. 1.

**Figure 6:**
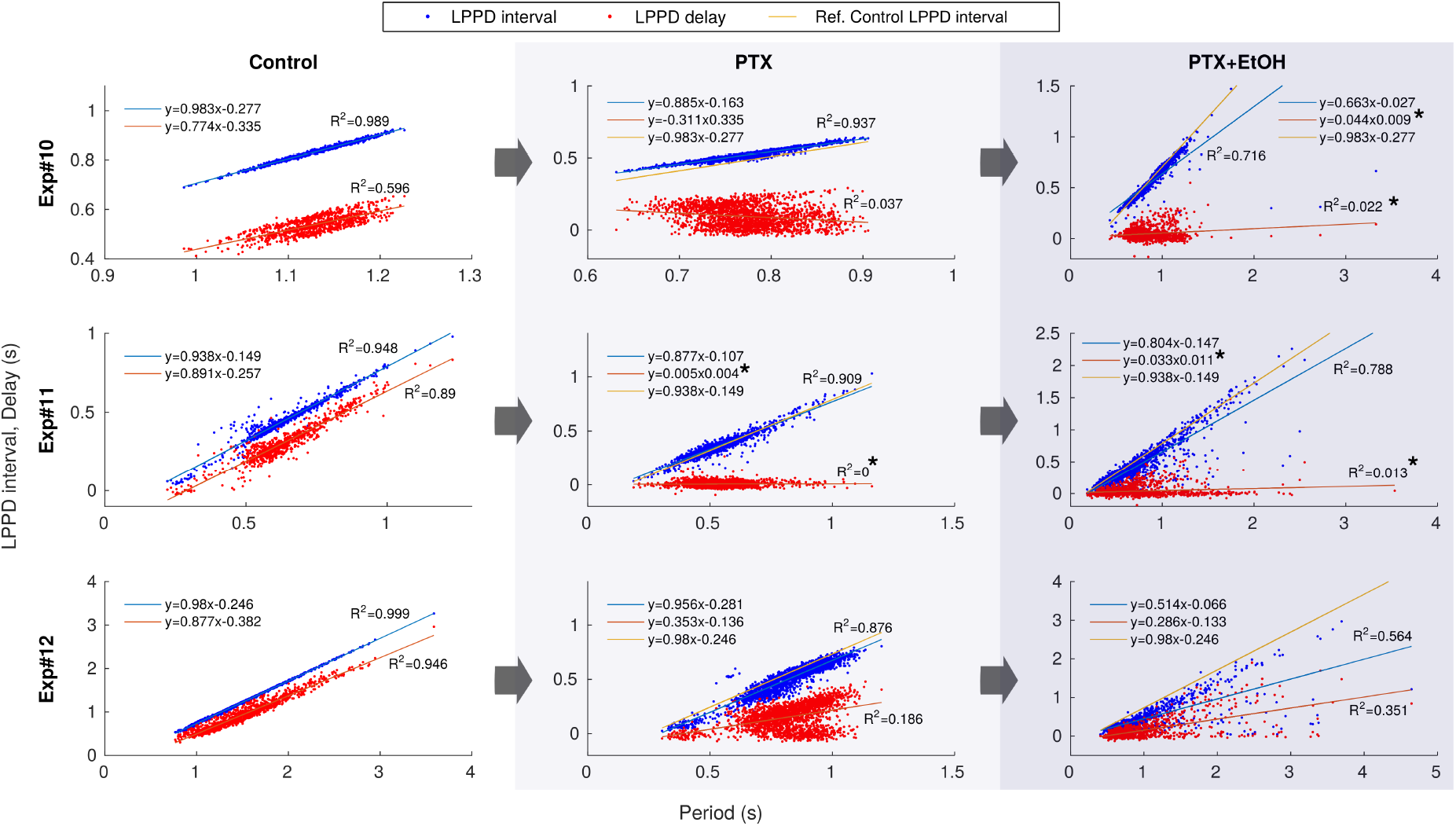
Comparison of the two dynamical invariants in three conditions: *control*, with *PTX* and with *PTX + Ethanol* in 3 different preparations (c.f. *Figure* 1). The correlation between the measured *LPPD interval* and *Period* is shown in blue while the correlation between the *LPPD delay* and *Period* is shown in red. Each point corresponds to one pyloric cycle. Regression analysis showed that both LPPD interval and delay values increased with period (*p* < 10^−7^ each). The linear dependence is indicated by *R*^2^ values displayed for each experiment in the corresponding panel. * Slope not significantly different from 0 (*p* > 10^−7^). Line in orange corresponds to the linear regression between the measured *LPPD interval* and *Period* in control conditions shown in the first column.

The presence of both invariants *LPPD interval*[*Period*] and *LPPD delay [Period]* is a very robust result, observed in all control experiments performed (*n*=42). In the supplementary material we provide a script to quantify and display the two reported invariants in any simultaneous recording of LP and PD neurons.

### 2.3 Cycle-by-cycle analysis

A major contribution of our work is the demonstration of cycle by cycle adjustments that give rise to the invariants, which can be indirectly be seen in ***Figure 3**, **Figure 4*** and ***Figure 6***. Even though in these figures each point corresponds to one pyloric cycle, the temporal relationship between points is lost in this type of representation. To better illustrate the instantaneous compensations of the different intervals in each cycle we highlight one representative example of transient changes in an intrinsically irregular preparation. This is shown in ***Figure 7*** (*Exp* #5 in ***Figure 3*** and *Figure* 4). Panel A in ***Figure 7*** shows the evolution of each interval *Period, BD_LP_*, *BD_PD_*, *LPPD delay, LPPD interval* and *PDLP delay* for each cycle period. One can observe that *LPPD delay, LPPD interval* closely follow the *Period* despite its variability. Panel B shows the similarity of each interval to the *Period* by subtracting each value from the period. Therefore, *LPPD delay*, *LPPD interval* remain now constant, since they change according to the *Period*, revealing both dynamical invariants. Note that the variability in the upper traces is only due to the Period variability (cf. *Period* trace in panel A). Panel C depicts all the intervals standardized so that the variability of all of them are presented in the same range. In this representation, the intervals that give rise to the invariants evolve on top of each other (see inset). However, the evolution of the intervals *BDLP*, *BDPD* and *PDLP delay* intertwine each other approximately compensating their variability among them. An example of this analysis under ethanol is shown in Supplementary Material.

**Figure 7:**
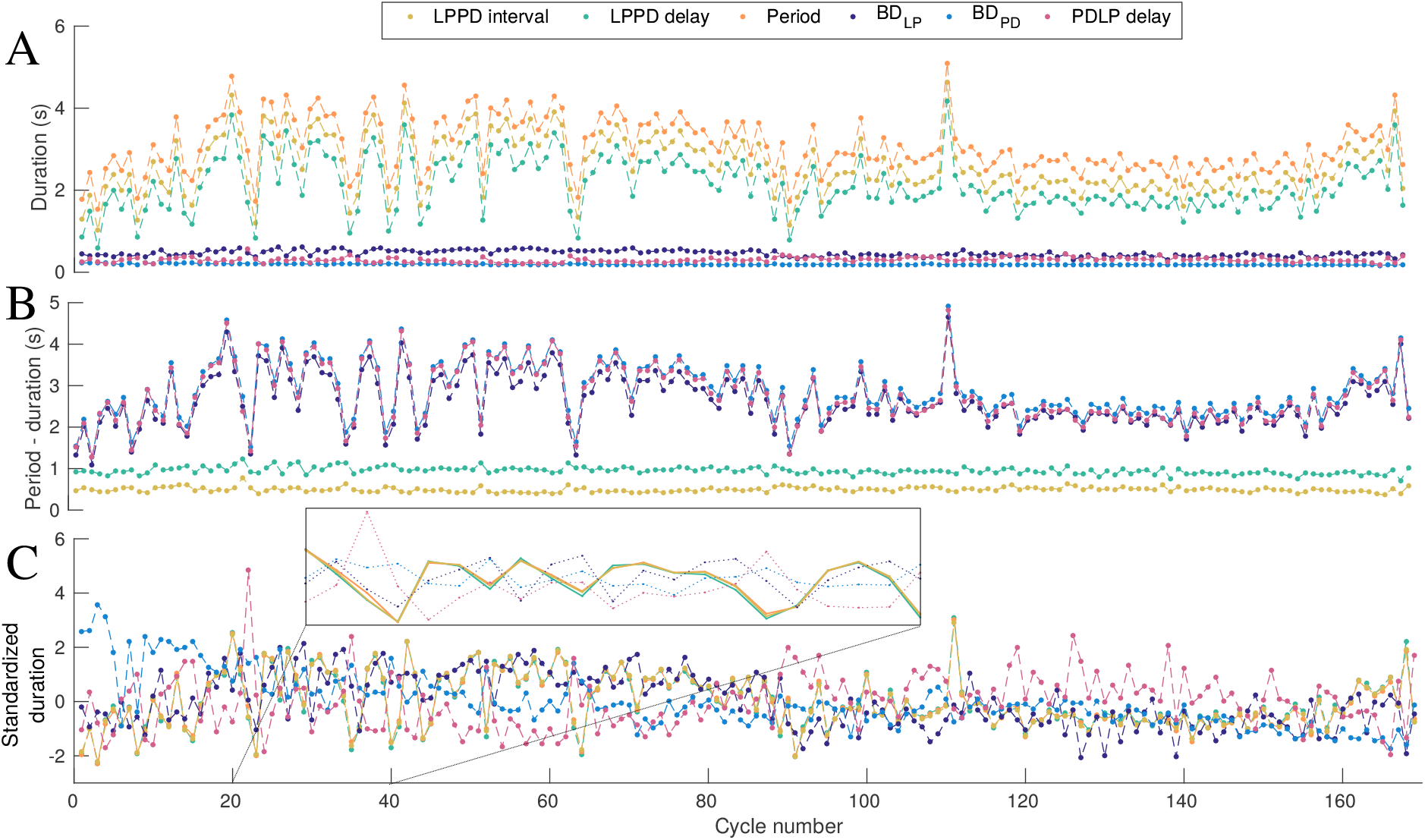
Cycle by cycle transient changes in the studied intervals. Panel A, intervals *Period, BD_LP_*, *BD_PD_*, *LPPD delay, LPPD interval* and *PDLP delay* for each cycle. Note that despite the variability in period, *LPPD delay* and *LPPD interval* closely follow it. Panel B shows the same intervals subtracted from the period. *LPPD interval* and *LPPD delay* associated intervals remain now constant, revealing both dynamical invariants. Panel C shows the intervals as in Panel A but with standardized duration. In this representation, the variability of all intervals are in the same range. Note that the standardized *LPPD delay*, *LPPD interval* and *Period*, which give rise to the invariants, evolve on top of each other while the evolution of the others intertwine. The inset shows a blow up to highlight the common evolution of the three intervals involved in the invariants (solid lines).

## 3 Discussion

Although typically characterized by frequency, most brain rhythms throughout the nervous system are built from sequential activations of groups of neurons.^12,58,59^ Some of these sequences are often very robust and directly related to the execution of motor commands, cognitive decisions and behavioral actions. When generating robust sequential activations, cycle-by-cycle flexibility and fine tuning of instantaneous periods, phases and event timings can be crucial for the optimization and the achievement of effective functions. In this paper, we have addressed this issue in a well-known experimental model where these questions can be more easily examined.

Using an adequate time reference frame and different experimental conditions to expose transient dynamics, our characterization of cycle-by-cycle variability in CPG circuits has revealed the presence of dynamical invariants in neural sequences. These invariants seem to arise from the rich intrinsic cell dynamics and the asymmetric connectivity balancing robustness and flexibility in the transients that build functional neural rhythms.

Most experimental and computational studies on CPGs focus their analysis on regular regimes, frequently discarding non-regular transient activity. However, the analysis of irregular CPG rhythms, rich in transient dynamics and not only in steady state activity, can unveil important properties of the robust neuron and network dynamics underlying rhythm coordination. Irregular rhythms were obtained in this study by two means: intrinsic irregularity, and biophysical disruption with ethanol. In particular, ethanol was very effective at revealing rich dynamics of neurons and connections in the pyloric circuit as seen by the larger flexibility of neuron activity. The effect of ethanol was reversible and in most cases the neurons returned to their original rhythm after ethanol was washed out or evaporated. Moderate ethanol application did not disrupt the anti-phase relationship between LP and PD neurons, thus the robustness of the sequence was kept, but evoked variable burst duration and hyperpolarization intervals.

As opposed to traditional regular activity recordings, irregular rhythms caused by intrinsic factors in the preparation presented high variability in hyperpolarization intervals and waveforms in both LP and PD neurons. LP presented larger plateaus and higher variability in burst duration while PD activity remained less variable. Irregularity induced by ethanol, however, presented remarkably flexible and long PD burst durations, while LP burst duration was more restricted. Ethanol also induced variability in the hyperpolarization intervals in both neurons. For further analysis, PTX was used along with ethanol to study how removing fast synapses in the circuit affected the invariants and tilted the interacting forces of the network that negotiate timings within a robust sequence.

### 3.1 Cycle-by-cycle analysis

Cycle-by-cycle analysis allowed to center the study of dynamical invariants in transient regimes, without losing the temporal relationship between key intervals building the sequence. Results show that *LPPD delay* and *LPPD interval* closely follow the changes in *Period* despite the variability underlying both dynamical invariants. The intervals, *BD_LP_*, *BD_PD_* and *PDLP delay* approximately compensate their variability and therefore contribute to the maintenance of the observed invariants.

The experimental results from the analysis of irregular CPG rhythms presented in this work illustrate that rich dynamics of neurons and connections in the pyloric circuit contribute to regulate flexibility and coordination to readily negotiate their specific timing within the sequential activity. The CPG tends to preserve cycle-by-cycle temporal relationships between neurons even under extreme conditions which points out to circuit’s highly effective negotiating properties and the dynamical arrangement of the motor rhythm to balance robustness and flexibility. CPGs could use the reported invariants and other preserved relationships to program their function under distinct circumstances, which may underlie their remarkable context-specific autonomous adaptability and functional efficiency.

### 3.2 Presence of dynamical invariants

In the sequential activity of the pyloric CPG, we identified the presence of cycle-by-cycle preserved relationships, and explored their role in the fast rhythm negotiating properties of this circuit. We found two dynamical invariants considering the measures analyzed: *LPPD delay* and *Period* and *LPPD interval* and *Period*. These invariants were present not only in regular control conditions but also in intrinsic irregular conditions and when high irregularity was induced by ethanol. In fact, the presence of both invariants *LPPD interval*[*Period*] and *LPPD delay*[*Period*] was a very robust result, especially in control conditions, in which the invariants were found in all experiments performed (*n*=42). One plausible explanation for the invariants is that variability in the PD and LP burst durations *BD_PD,LP_* and variability in the *LPPD delay* compensate each other in each pyloric cycle. The invariant *LPPD interval*[*Period*] is more precise than *LPPD delay*[*Period*], this is probably because *LPPD interval* contains the added variability of both *BDLP* and *LPPD delay*. It is important to note that other explored relationships among CPG activity intervals did not lead to invariants. This might imply that they are not so important for the rhythm negotiation and, thus, can remain constant or have unrelated variability with the period to fulfill another role. Since in our experiments we only used intracellular recordings of LP and PD neurons, we cannot discard the presence of additional preserved relationships among other neurons. Further study of dynamical invariants, including possible nonlinear relations, is required to fully understand their origin and implication in the generation of CPG rhythms.

It is important to emphasize that in this work we used different time intervals from the commonly used latency onset and offset, which are defined using the PD neuron first spike.^35,47,48,56^ When connectivity is asymmetric, the selection of the time references used to define the intervals is crucial for exposing potential dynamical invariants. LP neuron receives less connections than other neurons in the pyloric circuit. Therefore, it has more flexibility to adapt and coordinate its activity with the rest of the circuit elements in a cycle-by-cycle basis, making this neuron a better candidate as a time reference (see also ^48^).

We observed that the specific asymmetric connectivity of the pyloric network plays a key role in shaping the invariants. After removing glutamatergic synaptic inputs by applying PTX, the correlation *LPPD delay[Period]* was completely gone while *LPPD interval*[*Period*] was still maintained. Preservation of this last invariant is probably due to the LP burst duration variability, which manages to compensate the PD variability, even under the effect of ethanol.

### 3.3 Beyond CPGs

Most neural functions are supported by neuronal oscillatory activity, often simply referred to as a rhythm.^58,60^ Rhythms are recorded in specific brain circuits, such as in CPGs, or observed in recordings spanning distinct frequencies and anatomical regions, such as the cerebellum, the hippocampus, the basal ganglia and cortical areas. In most cases, brain rhythms are characterized and quantified regarding only their frequencies. However, a wide variety of experimental works show that robust sequential activations of different neuron types participate or are recruited at different phases of the oscillations that define brain rhythms (e.g.^61–66^).

Most neural rhythms, as pyloric neural oscillations, are based on inhibition as the main mechanism shaping not only the rhythmic activity,^67^ but most importantly, the sequential activation of its constituent elements. Inhibition based mechanisms offer specific time windows where neurons can express their excitability, balancing the robustness of the sequence and the flexibility to tune activation timings. Thus, asymmetric inhibition is a main player in creating and shaping functional sequences of cell activations. The actual execution of a sequential neural command, e.g., in the performance of a movement, is determined not only by the serial order of individual participants but also by their timing. This is the case for the pyloric CPG, as most likely fine timing adaptations are required to optimize the function of the motor plant beyond keeping the sequence needed to move food from one side to the other.

Thus, while executing a neural sequence, dynamical invariants might reflect important constraints to optimize functionality by shaping the actual intervals in which activity emerges in a coordinated manner. Here we have shown that the triphasic rhythm of the pyloric CPG displays a large degree of flexibility as quantified by the variability of the intervals that build the sequence. However, the dynamical invariants balance and constraint this flexibility in each cycle. It is important to note that the two invariants observed during cycle-by-cycle transients are different than the approximate average phase maintenance reported in previous works that used cross-preparations analysis, steady activity recordings, and other time references.^35,47,48,56^ Approximate phase maintenance, obtained in other works by averaging phase and periods in different preparations or in the same preparation under different treatments, might reflect some aspects of the unveiled invariants, but not their presence in cycle by cycle analysis, in particular during transients.

The unveiling of dynamical invariants in the spatio-temporal patterns of neural activity may have an important impact in robotics. Traditional robotic locomotion control paradigms are based on ad-hoc rules to deal with different scenarios (e.g. obstacle avoidance, uneven terrain, etc). The concept of dynamical invariants provides an alternative way to autonomously build constraints to drive behavior in all situations. In this context, a dynamical invariant based CPG control arising from the connectivity and rich intrinsic neuron dynamics^20^ can provide autonomous solutions to different situations informed by sensory feedback.

Beyond spiking-bursting activity and CPG function, dynamical invariants in other brain rhythms can underlie the creation of cyclic windows within oscillations when synaptic input can be most efficiently integrated for the effective execution of sequences generated in a given informational context. We foresee that the study of specific time references and dynamical invariants in different neural systems will provide novel views on the functional role of brain rhythms and their constituent sequences.

## 4 Methods and Materials

### 4.1 Electrophysiology

Adult male and female shore crabs (*Carcinus maenas*) were purchased locally and maintained in a tank with 13-15 C artificial seawater. Crabs were anesthetized by ice for 15 min before the dissection. The procedures followed the European Commission and Universidad Autónoma de Madrid, animal treatment guidelines. The stomatogastric nervous system was dissected following standard procedures and pinned in a Sylgard-coated dish containing *Carcinus maenas* saline (in *mM*: 433 *NaCl*, 12 *KCl*, 12 *CaCl_2_ · 2H_2_O*, 20 *MgCl_2_ · 6H_2_O*, 10 *HEPES*, adjusted to pH 7.60 with 4 *M NaOH*). After desheathing the STG, neurons were identified by their membrane potential waveforms and the spikes times in the corresponding motor nerves. Membrane potential from neurons was recorded using 3 *M KCl* filled microelectrodes (50 *M*Ω) and a DC amplifier (ELC-03M, NPI Electronic, Hauptstrasse, Tamm, Germany). Extracellular recordings were made using stainless steel electrodes in Vaseline wells on the motor nerve and amplified with an AC amplifier neuroprobe (model 1700, A-M Systems, Bellevue, WA, USA). Data was acquired at 10 KHz using a A/D board (PCI-MIO-16E-4, National Instruments). Spike timings were obtained from intracellular recordings in Dataview (https://www.st-andrews.ac.uk/~wjh/dataview/), using FIR filters and thresholds. Preparations were exposed to concentrations of (170 *mM*) Ethanol (Panreac), added directly to the bath. Glutamatergic synaptic inputs were blocked using 10^−7^ *M* picrotoxin (PTX; Sigma-Aldrich). Only preparations that completed all categories of treatment were included for each analysis.

### 4.2 Time references and interval measures

To further analyze and quantify these observations we have considered several measures based on precise time references at the beginning and at the end of the bursts (see***Figure 2*** left panel): PD and LP burst duration *BD_PD,LP_*: intervals from the first spike to the last spike of PD and LP neuron, respectively; *LPPD delay*: interval from the last LP spike to the first PD spike; *LPPD interval*: interval defined from the LP first spike to the PD first spike; *PDLP delay*: interval from the last PD spike to the first LP spike in the following burst; *Period:* interval from first LP spike to the next first spike in the following LP burst. We quantified these measures in long intracellular recordings (15 min on average). There were some extreme cases in irregular rhythms induced by ethanol where time references were not well defined and the corresponding activity had to be removed from the statistics shown below. An example of these cases is illustrated in ***Suplementary figure* 3**. The number of bursts that had to be dismissed in these experiments ranged from 1 to 17% and 0 to 27% of the total number of bursts of LP and PD neurons, respectively, in the recordings (see ***Suplementary table* 1**).

The coefficient of variation defined as *C_v_* = *σ/μ* · 100 depicted in the boxplots ***Figure 2*** and ***Figure 5*** was calculated as an average of the *C_v_i__* of each experiment in an ensemble *n* specified in each plot.

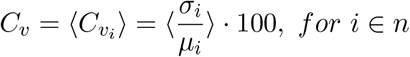

The calculated P-values clustered clearly in two groups; higher values ranged from 0.8 to 10^−7^, while smaller values ranged from 0 to 10^−16^. Thus, the significance level *α* used for the null hypothesis significance test for correlations was *α*= 10^−7^.

Standardized cycle intervals *z* in ***Figure 7*** were calculated as follows:

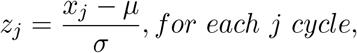

where *x* is the interval value.

Experimental data analysis was implemented with Matlab, the scripts are available in the supplementary material, including those for the invariant graphic representations. These scripts can be used for further validation in other CPG circuits and, in fact in any other candidate neural sequence.

The video in the supplementary material was rendered using the python library *matplotlib* and the audio was processed using Dataview sonification tool.

## 5 Acknowledgments

We thank Manuel Reyes-Sanchez and Carlos Garcia Saura for assistance with the video. This work has been supported by Spanish grants MINECO DPI2015-65833-P, TIN2014-54580-R, TIN2017-84452-R (http://www.mineco.gob.es/), and ONRG grant N62909-14-1-N279.

## 6 Author contributions

I.E., R.L. and P.V. conceived and designed the experiments. I.E. and R.L. performed the experiments. I.E., R.L., D.A., F.B.R. and P.V. analyzed the data. I.E., R.L. F.B.R. and P.V. wrote the paper.

## Supplementary Material

### Figures and tables

**Supplementary Figure 1:**
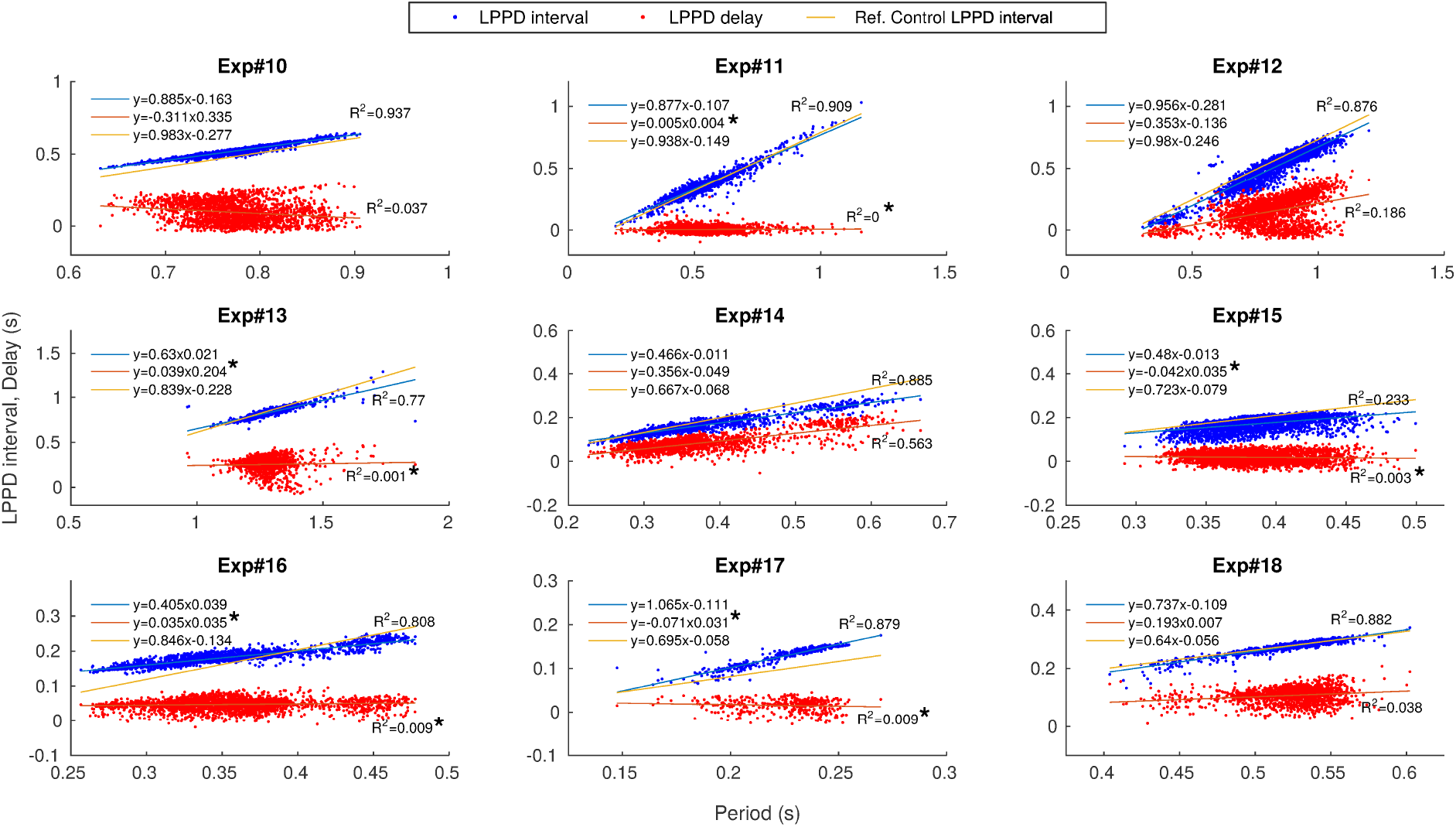
Comparison of the two dynamical invariants after applying PTX 5 · 10^−7^ *M* in 9 preparations. The correlation between the measured *LPPD interval* and *Period* is shown in blue while the correlation between *LPPD delay* and *Period* is shown in red. Each point corresponds to one pyloric cycle. Linear regression is depicted for each experiment. Regression analysis showed that both LPPD interval and delay values increased with period (*p* < 10^−7^ each). The linear dependence is indicated by *R*^2^ values displayed for each experiment in the corresponding panel. * Slope not significantly different from 0 (*p* > 10^−7^). Line in orange corresponds to the linear regression between the measured *LPPD interval* and *Period* in control conditions.

**Supplementary Figure 2:**
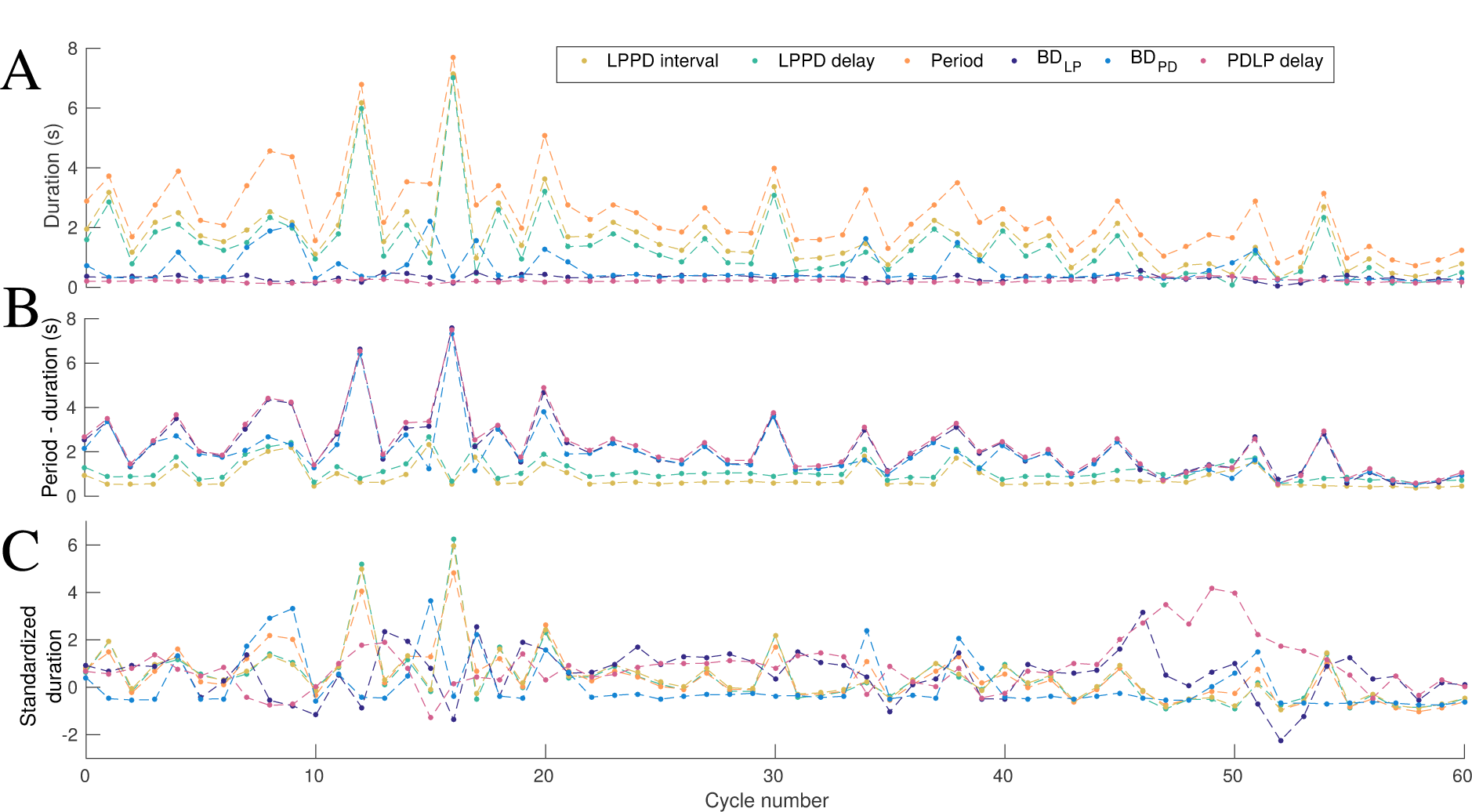
Cycle by cycle transient changes in intervals under ethanol. Panel A: intervals *Period, BD_LP_*, *BD_PD_*, *LPPD delay, LPPD interval* and *PDLP delay* for each cycle. Note, that period is variable and *LPPD delay* and *LPPD interval* closely track it. Panel B shows the same intervals subtracted from period. The interval that are less variable, i.e., *BD_LP_*, *BD_PD_*, do not follow the change in period. Panel C shows the same intervals in A but with standardized duration.

**Supplementary Figure 3:**
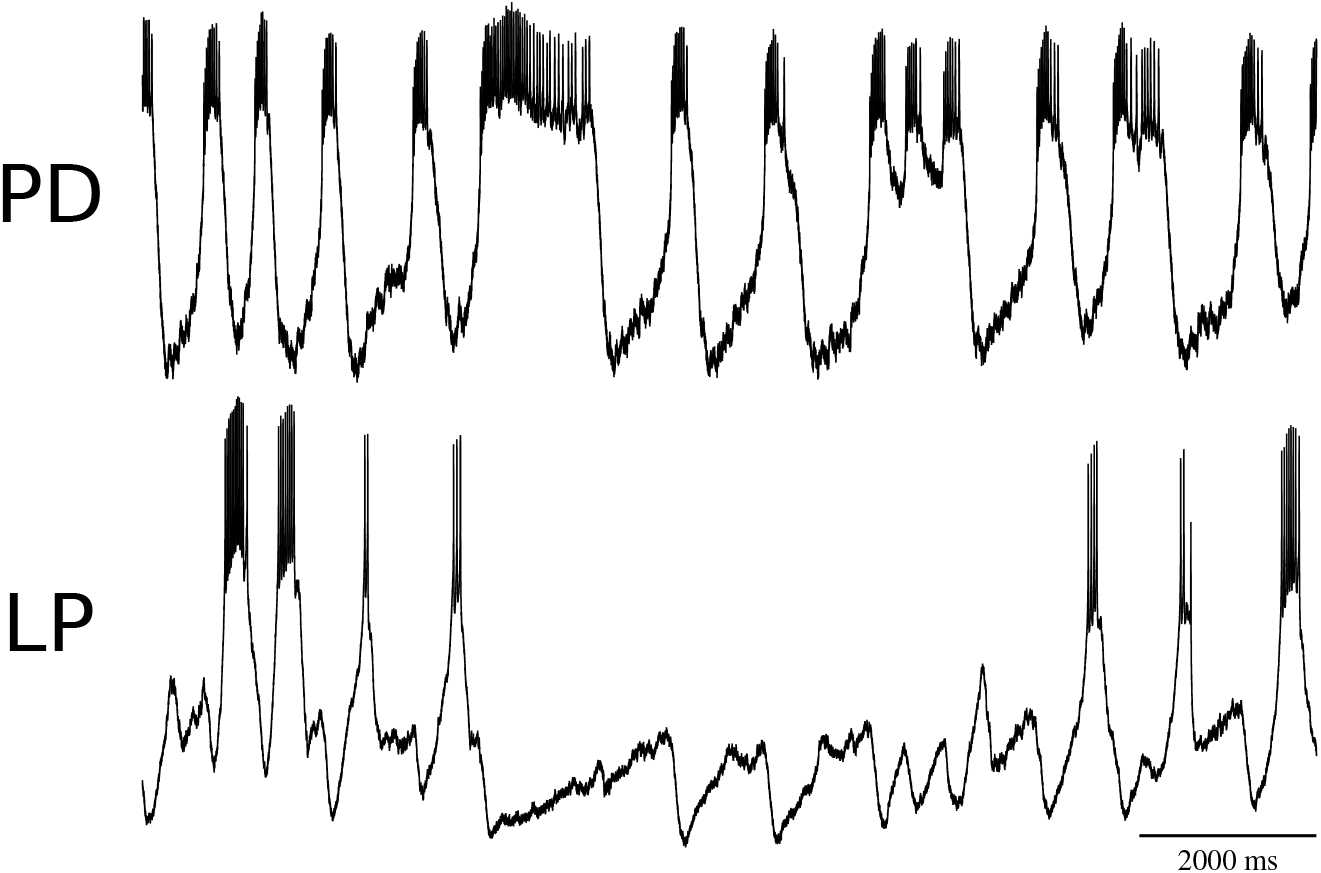
Example of highly irregular rhythms in which the measures used to characterize the rhythm and its variability (LP, PD burst durations, LPPD last-first period and first-first LP period) cannot be defined and the corresponding activity had to be dismissed in our analysis because the time references used are lost,. LP neuron stops bursting while PD sustains its activity.

**Supplementary Table. 1:** Percentage of dismissed bursts of the total number of burst of LP and PD neurons in all experiments because of missing time references (see ***Suplementary figure* 3**).

| # Experiment | Dismissed LP bursts | Dismissed PD bursts |
| --- | --- | --- |
| 1 | 2% | 6% |
| 2 | 3% | 2% |
| 3 | 4% | 11% |
| 4 | 12% | 13% |
| 5 | 5% | 4% |
| 6 | 2% | 1% |
| 7 | 17% | 26% |
| 8 | 10% | 27% |
| 9 | 10% | 12% |
| 10 | 2% | 3% |
| 11 | 2% | 4% |
| 12 | 7% | 4% |
| 13 | 1% | 1% |
| 14 | 6% | 16% |
| 15 | 3% | 1% |
| 16 | 2% | 2% |

### Video 1

Video of the evolution of the time intervals giving rise to dynamical invariants. Left panel: LP and PD voltage time series showing the instantaneous *LPPD interval* (blue), *LPPD delay* (red) and *Period*(black) in an illustrative experiment. Right panel: Evolution of the dynamical invariants in time along the regression lines. Stereo sound corresponds to the sonification of LP (left ear) and PD (right ear) neurons which helps to detect the rhythm variability. Video summarizes a 27 min recording extracting interesting transitions with rhythm accelerations and decelerations that entail a sudden change in *LPPD interval*[*Period*] and *LPPD delay*[*Period*].

### Matlab scripts

Matlab scripts that calculate the intervals defined in the text from the spike-times and plot the invariants and barplots of the coefficient of variation. These scripts can be used for further validation in other CPG circuits and, in fact, in any other candidate neural sequence.

